# Increased reliance on temporal coding when target sound is softer than the background

**DOI:** 10.1101/2023.06.19.545590

**Authors:** Nima Alamatsaz, Merri J. Rosen, Antje Ihlefeld

## Abstract

Everyday environments often contain multiple concurrent sound sources that fluctuate over time. Normally hearing listeners can benefit from high signal-to-noise ratios (SNRs) in energetic dips of temporally fluctuating background sound, a phenomenon called dip-listening. Specialized mechanisms of dip-listening exist across the entire auditory pathway. Both the instantaneous fluctuating and the long-term overall SNR shape dip-listening. An unresolved issue regarding cortical mechanisms of dip-listening is how target perception remains invariant to overall SNR, specifically, across different tone levels with an ongoing fluctuating masker. Equivalent target detection over both positive and negative overall SNRs (SNR invariance) is reliably achieved in highly-trained listeners. Dip-listening is correlated with the ability to resolve temporal fine structure, which involves temporally-varying spike patterns. Thus the current work tests the hypothesis that at negative SNRs, neuronal readout mechanisms need to increasingly rely on decoding strategies based on temporal spike patterns, as opposed to spike count. Recordings from chronically implanted electrode arrays in core auditory cortex of trained and awake Mongolian gerbils that are engaged in a tone detection task in 10 Hz amplitude-modulated background sound reveal that rate-based decoding is not SNR-invariant, whereas temporal coding is informative at both negative and positive SNRs.

## Introduction

A signature problem for most hearing-impaired individuals is their reduced ability to dissociate target from background sound. When an auditory target and a background sound source coincide, the background sound can energetically mask the target by swamping or occluding the target’s cochlear representation, a problem that even ideal rehabilitative listening devices cannot solve. However, because natural sounds inherently fluctuate over time, they rarely overlap continuously [1]. To segregate competing sound sources into perceived auditory objects, both humans and animals can exploit these temporal fluctuations [2, 3, 4], a phenomenon called “dip-listening.” To best define treatment targets for hearing loss, it is necessary to identify coding strategies for dip-listening wherein behavioral ability to detect a tone in temporally fluctuating background sound co-varies with target-evoked neuronal activation.

Neurophysiological studies in animal models have greatly advanced our mechanistic understanding of dip-listening. Specialized neuronal rate- and temporal coding mechanisms enhance target representations during high-and-short-term peaks in SNR across multiple processing stages, including cochlea [5], cochlear nucleus [6], inferior colliculus [7] and auditory cortex (ACx, [8]). In addition, dip-listening can be behaviorally quantified by comparing detection, discrimination or identification thresholds between temporally fluctuating vs. steady maskers. The improvement in thresholds, called modulation masking release (MMR), is refractory to training and varies with SNR. At positive SNRs, where target energy dominates the acoustic mixture, MMR is not observed across a range of tasks. Thus for positive SNRs, a coding strategy based on net change in long-term acoustic energy (rather than on rapid temporal fluctuations) may suffice to detect a target sound. However, at negative SNRs, where dip-listening can dissociate the target from the acoustic mixture if the masking envelope fluctuates moderately (4-32 Hz, peaking around 10 Hz), MMR increases with decreasing SNR [9, 10]. This suggests that SNR shapes the reliance on short-term temporal processing, thereby modulating how listening in fluctuating background sound operates. To test this hypothesis, we here study ACx because this is where auditory objects emerge, making ACx a promising target while searching for mechanisms allowing dip-listening to operate across a range of negative and positive SNRs. Moreover, a mechanism thought to underlie dip-listening, envelope locking suppression, reduces the fidelity by which ACx neurons track background sound when a target occurs, at both positive and negative SNRs, at least in anesthetized animals [11, 7]. However, a mechanistic understanding of how SNR shapes the ACx neuronal code is complicated by the fact that prior neurophysiological work assessing dip-listening often uses untrained animals or animals that could not detect target sound at negative SNRs [12, 13]. Here we are interested in SNR invariance of auditory cortical responses in awake gerbil during dip-listening. Gerbils have low-frequency sensitivity and MMR of comparable magnitude as humans [14, 15], making them a suitable model for studying SNR invariance during dip-listening.

Using appetitive psychometric testing and chronically implanted recording electrodes, we simultaneously quantify behavioral sensitivity and ACx single-unit responses. Awake trained animals either actively detect a tone in modulated masking noise, at three SNRs (−10, 0 and 10 dB), or passively hear the same sounds without task engagement. Behavioral accuracy is comparable across SNRs, presumably controlling for task difficulty. In experiment 1, we contrast the effect of SNR on rate vs. temporal coding strategies in sound-detecting animals. We previously discovered that a measure of discriminability between the sound-evoked cortical responses when target sound is present vs. absent, mutual information based on firing rate, is smaller for behaviorally relevant targets as compared to sound of no behavioral significance, hinting that behavioral relevance may increase reliance on temporal as opposed to rate coding [16]. To elucidate the role of behavioral relevance, in experiment 2 we test non-sound-detecting animals, using the same sounds and similar operant conditioning as in experiment 1 (except for giving response reward irrespective of decision). Our results support the interpretation that temporal coding can directly serve to detect target sound across all tested SNRs, whereas the neuronal information carried in rate coding needs a more nuanced readout strategy with differing decision criteria at positive vs. negative SNRs.

## Materials and Methods

All experimental protocols were approved by the Rutgers University Institutional Animal Care and Use Committee.

### Housing

Animals were group housed unless warranted by veterinary exemption for post-surgical recovery. Environmental enrichment was provided with soft nesting materials and toys for chewing. The gerbils had unrestricted access to a nutritionally complete diet through food pellets along with unrestricted water prior to training and during water breaks, as well as controlled water during testing. Life-brand cereal was given as treats after each training session. For chronically implanted animals, dietary supplements such as diet gel, hydrogel, and sunflower seeds were provided before and after the surgical procedure. Daily checkups ensured animal health and safety. The vivarium temperature (65-75°F) and humidity (< 50%) were logged and maintained, and the vivarium lights were automatically switched throughout the day to regulate sleep-wake cycle of the animals.

### Behavioral Testing

Gerbils were placed inside a testing cage with a nose poke to initiate or abort a trial and a lick spout to access water (Figure 1A). The entire setup was inside a radio frequency shielded sound-attenuating booth with sound-absorption walls (booth dimensions: 8’ x 10’ x 7’). A loudspeaker was mounted on the booth ceiling, approximately 1 m above the center of the cage.

**Figure 1.**
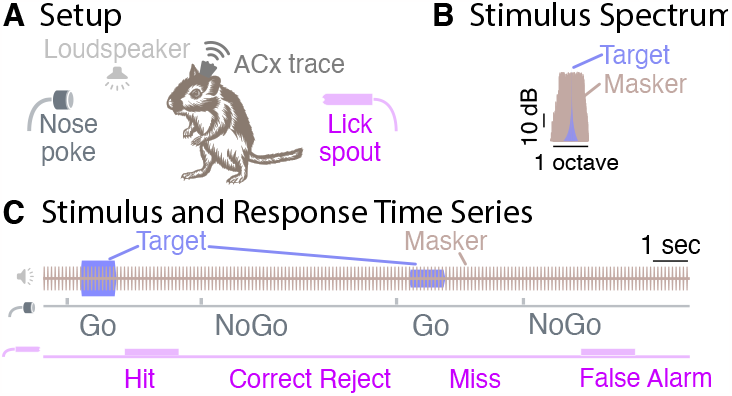
Testing apparatus and behavioral design. **A** The test setup included a loudspeaker above the test area, a nose poke and a lick spout. In addition, for chronically implanted animals, a wireless system recorded the cortical traces. **B** The background sound (brown), consisting of 10 Hz amplitude modulated noise, was continuously present. On Go trials, a target sound (blue) was additionally played, consisting of a 1 s, 1 kHz tone and randomly chosen from -10, 0 or 10 dB SNR. **C** The gerbil triggered a new trial by breaking a light beam inside the nose poke, and could obtain water reward through the lick spout. A loudspeaker mounted above the test area played the sounds. The gerbil then responded to the trial condition either by licking the water spout, or by withholding a response through waiting or by poking the nose poke once more. Depending on the stimulus condition, this response resulted in either a Hit, a Correct Reject, a Miss or a False Alarm.

For experiment 1, using an appetitive Go/NoGo paradigm with controlled water access (Figure 1C), six adult Mongolian gerbils (*Meriones unguiculatus*) were trained and tested on a tone detection task while fluctuating noise, called *masker*, continuously played in the background throughout each session [15]. Specifically, gerbils detected whether or not a 1 kHz target tone (1 s duration, 50 ms cosine-squared rise/fall, 40-60 dB SPL) was present in 10 Hz rectangularly amplitude-modulated and band-limited background noise (50% duty cycle, 10 ms cosine-squared rise/fall ramp on each rectangular burst, 50 dB SPL, 2/3 octave bandwidth, pseudorandom frozen noise, centered at the target frequency of 1 kHz; Figure 1B).

The sequence and timing of training and testing is depicted in Figure 2. Gerbils were initially trained to associate the 1 kHz, 60 dB SPL target tone with water, then learned to use a nose poke to initiate a target tone (*Go* trial) and finally were taught that on some trials, despite nose poking, no target tone would play and no water reward would be given (*NoGo* trial). Specifically, to initiate a trial, gerbils were trained to hold their nose inside the nose poke for 200 ms and wait for a target stimulus to potentially occur for a potential water reward from the lick spout.

**Figure 2.**
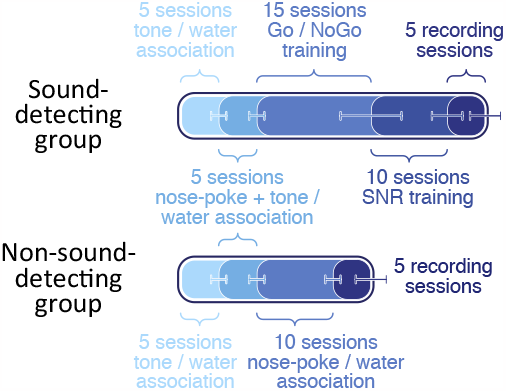
Average number of sessions per each training and testing stage shown as shaded progress bars for each of the two gerbil groups. The lower and upper bounds of session counts are indicated with error bars.

On Go trials, a target was presented in the continuous background sound, whereas on NoGo trials, no target was played and thus only the background could be heard (compare light blue target vs. light brown masking sound traces in Figure 1C). On Go trials, gerbils were then supposed to approach a water spout within a 4.2 s period after poke onset for a water reward. Successful contact with the spout during Go trials was scored as HIT. Other responses (either a re-poke in the nose poke or no poke response) were scored as MISS. On NoGo trials, gerbils were supposed to withhold any lick spout response. A withheld lick spout response or a re-poke response to the nose poke were each scored as CORRECT REJECT. However, if gerbils approached the water spout during a NoGo trial, no water was released, a 1-1.5 s mandatory timeout was given and the response was scored as FALSE ALARM (compare light blue / brown sound, grey nose poke and blue lick spout traces in Figure 1C). To discourage guessing, each FALSE ALARM trial was immediately followed by another NoGo trial for a maximum of 15 sequential NoGo trials. For each session, the rate of hit responses (HR) and the rate of false alarms (FAR) was then used to calculate the behavioral sensitivity, called d,, according to equation 1:

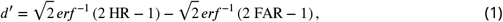

where *erf* ^−1^ is the inverse error function. Test sessions typically lasted 60 minutes, but varied in duration, depending on the animal’s satiety and willingness to perform the task. Averaged across sessions and animals, animals typically performed approximately 108 Go trials and 62 NoGo trials per session. To computationally eliminate the possibility of infinite d,, both (HR) and (FAR) were conservatively bracketed via thresholding such that any observed rate below 0.05 was assumed to equal 0.05, and any rate exceeding 0.95 was assumed to equal 0.95. Those thresholds were chosen assuming one guessed trial per SNR per session.

For both training and testing, the masker was fixed throughout each session. To ensure a fixed phase delay between target and masker, after initiation of a trial via nose poke, the target onset was delayed until the next 45° phase of the masker occurred. As a result, the target onset could occur between 250 to 350 ms after onset of a nose poke. This time delay arose as the sum of the 200 ms hold duration for the nose poke, a fixed 50 ms delay imposed due to limitations of the recording system for updating the acoustic output, and a 0 to 100 ms phase delay, depending on the phase when the animal initiated the nose poke during the 100 ms masker cycle.

All animals were initially trained with a masker level of 20 dB SPL and a target level of 75 dB SPL (55 dB SNR), until they reached criterion performance (FAR below 30% and d, above 1.5). Across sequential sessions, the masker was then gradually raised in steps of 10 dB, to a final masker level of 50 dB SPL (25 dB SNR). In the final training stage, target sounds at additional, softer intensities were gradually added in 5 or 10 dB steps across sessions, resulting in a total of three SNRs. The experimenter’s decisions to increase the masker or decrease the target levels was guided by whether or not the animal reached criterion performance at all tested SNRs and how confident the animal appeared in the task. All gerbils completed training within 30 sessions or less. At the conclusion of training, all gerbils were able to reliably detect tones at -10 dB, 0 and 10 dB SNR.

For testing, the masking noise was continuously present in the background at 50 dB SPL. Three target intensities (40, 50 and 60 dB SPL) were randomly interleaved from trial to trial, resulting in SNRs of -10 dB, 0 dB and 10 dB. In addition NoGo trials, without target energy, were randomly interleaved (SNR = −oo dB) After animals completed at least three sessions with d, of 1.9 or better at all SNRs, four of these sound-detecting gerbils advanced to the neural recording stage.

The two other sound-detecting gerbils were advanced to a separate behavioral test, to measure the minimum tone duration needed to perform the tone detection task. These two gerbils were behaviorally tested on tone detection at 0 dB SNR (50 dB SPL target level and 50 dB SPL masker level) as a function of target tone duration, across 13 sessions. Specifically, in each test session, seven different target tone durations (50, 100, 200, 400, 600, 800, 1000 ms) were randomly interleaved from trial to trial and the gerbil’s behavioral responses were recorded.

To control for the effects of arousal in the neural recordings, two additional gerbils were trained for experiment 2, using a non-sound-detecting Go/NoGo paradigm (Figure 2). Similar to the sound-detecting gerbils in experiment 1, these two gerbils were initially trained to associate the target tone with water, and then to trigger target tones by using the nose poke. However, unlike the sound-detecting gerbils, the non-sound-detecting gerbils then moved to a different training stage where water reward was only contingent on initiating a trial via nose poke and on reaching the water spout on time. Therefore, both Go trials (when target and background sound was played) and NoGo trials (when only the background sound was played) could result in water rewards. Once these two animals reached a minimum 95% (HR) and (FAR), indicating that they did not behaviorally discriminate between Go and NoGo before approaching the lick spout, they were advanced to the neural recording stage.

### Surgical Procedure

For the neural recording stage, we chronically implanted recording electrodes into left ACx [17, 18, 19]. Specifically, gerbils were initially anesthetized with 4% isoflurane, 1.5 mg/kg ketoprofen, 0.35 mg/kg dexamethasone, and continuously given 1-3% isoflurane to maintain sedation. Using stereotactic coordinates, with the medial-rostral corner at 5 mm lateral and 5 mm rostral to the *λ* landmark, a craniotomy was performed (approximate extent 1mm x 2mm). Using a 25 degree approach angle relative to the vertical axis, the targeted implantation site was then ascertained at 4.6 lateral and 3.4 mm rostral to *λ*. At the targeted site, a durotomy was performed before a silicon microelectrode array with 16 channels (A4×4-4mm-200-200-1250-H16-21mm, NeuroNexus Technologies, Inc.) was lowered into the brain at an initial insertion depth of 1.3-1.5 mm from the surface of the brain. A custom-made microdrive held the array in place inside a recording chamber, enabling post surgical advancement of electrodes, deeper into the brain tissue. The head-post along with the microdrive and implanted array were fixed on the skull using 4-6 bone screws and two layers of dental acrylic.

### Recording System

Trained gerbils were tested while cortical potentials were simultaneously recorded, amplified and transmitted wirelessly to a receiver positioned approximately 1 m from the cage (W16, Triangle BioSystems International). All input/output channels were synchronized at 100,000 samples/s and 16 bits by sharing the sampling clock pulse of two data acquisition cards (DAQs, PCIe-6321 and PCIe-6341, National Instruments Corporation) via a Real-Time System Integration bus cable. Specifically, custom written software, called Electrophysiology Auditory Recording System (EARS), synchronously controlled auditory stimuli delivery and recorded both behavioral and physiological responses [20]. EARS communicated with the loudspeaker, nose poke, licks spout, W16 and a personal computer for data storage and analysis via the two DAQs. In addition, EARS interfaced with a syringe pump (NE-1000 Programmable Single Syringe Pump, New Era Pump Systems, Inc.) via a USB-RS232 emulator.

#### Audio Delivery

Auditory stimuli were generated in EARS, D/A converted at a sampling rate of 100kHz and 16 bits, and preamplified (E 12:2, Lab.gruppen) before being sent to the loudspeaker (DX25TG59-04 tweeter, Tymphany HK Ltd). Sound calibration was initially performed by playing 18 bits long maximum length sequence (MLS) twice in a row, recording the response with a sound level meter placed in the center of the cage (Brüel & Kjær 2250), and inverting the cross-correlation between the second portion of the MLS and microphone recording to generate a pre-amplification audio filter that flattened the speaker response. Periodic re-calibration verified acoustic integrity of the recording system with +/-2 dB precision from 0.5 to 8 kHz.

#### Data Acquisition

Furthermore, custom printed circuit boards, connected to nose poke and lick spout, drove infrared emitter diodes (LTE-302, LITE-ON Technology Corporation) and their paired photosensors (OPS693, TT Electronics Plc), and conditioned these optical channel responses before sending them to their appropriate DAQ digital input channels. In addition, 15 analog input channels collected cortical potentials from the wireless receiver at a sampling frequency of 31.25 kS/s per channel.

### Behavioral-Cortical Assessment

On each recording day, cortical activity in animals was recorded twice, once during active task engagement mode, and later while the animal passively heard similar stimuli in a randomly different order of presentation. At the end of each recording session, electrodes were manually advanced by turning an advancing screw on the custom-made microdrive by approximately 40 *μm*.

### Analysis

The data analysis pipeline is depicted in Supplemental Information (Figure 1). Raw recorded cortical traces along with their associated behavioral events were analyzed offline in MATLAB (R2017a, The MathWorks Inc), by subtracting from each recording channel the grand mean across all channels and band-pass filtering this bias-corrected trace (zero-phase Butterworth, 6th order, 300-6000 Hz). Next, the resulting band-limited trace were time-windowed around the response time interval. Specifically, response time intervals were defined as starting at -1 s before potential target onset and ending +1 s after potential target offset, with the next 45° phase of the masker after the nose poke onset denoted as 0 s. In other words, 0 s marks the time point when the acoustic target onset occurred during Go trials, or the time when the target would have started during NoGo trials. Next, spike events were detected as negative peaks exceeding a threshold of 4.8*0*_*n*_, where *0*_*n*_ was the estimated standard deviation of the background noise [21]. All events with amplitudes exceeding 30*0*_*n*_ were rejected as artifacts.

The extracted event waveforms were further processed into putative units with an automatic spike sorting algorithm, using Principal Component Analysis (PCA) and k-means clustering algorithm (UltraMegaSort 2000, [22]). Visual inspection of each putative unit then verified that the shape of the unit’s waveform conformed to a time series typical for an action potential, that the firing rate of this unit was stable throughout the session, that there were infrequent refractory period violations (less than 1%), and that each putative unit was separated in at least one principal component space plot or with the best linear discriminant.

### Statistical Comparisons

Units that fulfilled all criteria were labelled as single-unit and further analyzed using seven rate and temporal coding metrics defined below. To assess whether rate vs. temporal response codes varied with SNR and across active and passive listening conditions, repeated measures or mixed design analysis of variance (rANOVA or mANOVA) was implemented with the *rstatix* package version [23] in R version 4.0.3 [24]. A significance level of *α* = 0.001 was consistently used throughout the analysis. Whenever the sphericity assumption for within-subjects factors was violated (based on Mauchly’s test), Greenhouse-Geisser correction was applied to the results. In addition, behavioral psychometric functions of (HR) were estimated using generalized linear models (GLM, [24]), assuming binomial response distributions (link function: logit), and weighing HR proportionally to the trial count of each stimulus condition.

### Response Time Histograms

For each unit and SNR during Go trials, as well as for NoGo trials, response time histograms (RTHs) were calculated with 50 ms resolution, by first binning spike events into 10 ms time-windows and then convolving the resulting response probability densities with a 50 ms rectangular kernel. This resolution was chosen because the slowest time-window to capture envelope-related acoustic responses is 50 ms, corresponding to the Nyquist rate of the 10 Hz masking envelope.

### Firing Rate

For each single-unit and SNR, firing rates were calculated by counting the number of spike events during incremental time-windows in steps of 50 ms starting at the target onset, and dividing it by the length of the window in seconds. The across-trial average and standard deviation of the resulting firing rates were then used to calculate the separation between Go and NoGo responses. Specifically, for each SNR, neurometric rate z-scores were calculated as a function of time, according to equation 2:

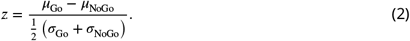

Specifically, for the rate z-score, *μ*_Go_ and *0*_Go_ were the average firing rate and standard deviation of firing across Go trials for each target SNR, and *μ*_NoGo_ and *0*_NoGo_ were the average and standard deviation of firing rate for the NoGo trials.

### Power Spectral Density

To estimate the spectral density of the single-unit responses, point process multi-taper spectrum (MTS) analysis was derived from the spike event times and pooled across trials [25]. Sparse single-unit events impede reliable MTS estimates at the single-trial level, a limitation solved here by sampling, for each SNR, half of the trials randomly with replacement, across 20 repetitions, using equation 2.

### Vector Strength

To calculate the strength by which single-units responded at a consistent phase of the masker envelope for each individual trial, using equation 3, we calculated the vector strength (VS) at 10 Hz.

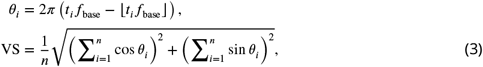

where *t*_*i*_ is the spike time relative to target onset, *f*_base_ is the base frequency at which VS is calculated (ranging from 1 to 20 Hz in steps of 1 Hz), *0*_*i*_ is the phase of each spike, and *n* is the total number of spikes (across trials, per target level). Using equation 4 to approximate the p-value for Rayleigh’s test for uniformity [26], across all NoGo trials and for each SNR during the Go trials, we then ascertained whether the VS to the masking envelope differed from zero.

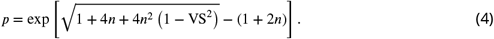

### Target-Evoked Decorrelation Response

To quantify how much the temporal response pattern of single-unit responses changed when a target tone was added to the background sound vs. when just the background sound was present, for each SNR and single-unit, we compared RTHs across Go vs. NoGo trials. Specifically, Pearson’s correlation coeffcient *ρ*, between Go and NoGo RTHs were calculated in running time-windows of 300 ms duration, sliding across the full duration of the response interval from -1 s to 2 s, with a step size of 10 ms. The 300 ms duration was chosen conservatively based on the results of the behavioral tone duration thresholds (see Figure 4). A high *ρ* indicates a high trial-to-trial similarity in the single-unit RTH between the masker-only response during NoGo trials and the target-and-masker-combined response during Go trials. In addition, target-evoked decorrelation responses were derived by calculating the *ρ* z-score between the masker-only RTH immediately preceding the target onset vs. the early response during the response time-window (equation 5, where *μ*_onset_ and *0*_onset_ were the average and standard deviation of *ρ* during the first 300 ms of target response, and *μ*_poke_ and *0*_poke_ were the average and standard deviation of *ρ* within 300 ms of the nose poke events).

**Figure 3.**
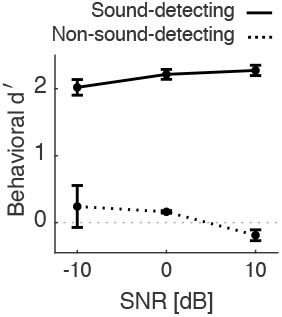
Average behavioral performance of each group of gerbils across the recorded sessions with active engagement (sound-detecting n=4, non-sound-detecting n=2, 5 sessions each). Error bars show one SEM.

**Figure 4.**
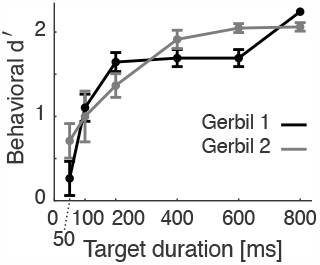
Behavioral curves at 0 dB SNR as a function of tone duration for non-implanted gerbils (n=2, sessions=13 each on average). Error bars show one SEM.

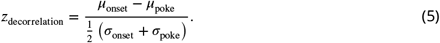

### Mutual Information

Under the assumption that during the sustained response interval (the interval from 0.1 s to 0.95 s), neural firing follows a stationary independent identically distributed Poisson process, we estimated *λ* parameters of the underlying Poisson distributions. *λ* was used to approximate mutual information [16] as an agnostic measure for comparing response similarity. Mutual information ranges from 0 to 1, with higher values indicating low similarity between the Go and NoGo response, and hence, higher transfer of information (yielding higher discriminability) over the target-evoked response channel. Conversely, lower mutual information can be interpreted as sparsity of the neural code in conveying information. Because mutual information was calculated on spike probability distributions, this is a rate-based neurometric.

### Similarity Index

To quantify the overall similarity of the sound-evoked responses to the mixture of target and background sound vs. just the background sound, for each single-unit, a similarity index was calculated using a metric defined in prior work [7]. Specifically, for each SNR, the time series of each unit’s Go and NoGo RTHs during the sustained response interval were plotted against each other and the resulting scatter plots were fitted using linear regression. The slope of these regression lines is called similarity index, and combines both rate and temporal information [7]. A similarity index of 1 indicates that when a target occurs, the response to the mixture of target and background sound is similar to the response to just the background sound, whereas a zero similarity index indicates that the presence of target energy strongly changes how a unit responds.

## Results

### Sound-Detecting Gerbils

Across all SNRs, the four implanted sound-detecting gerbils could reliably detect the target tone (solid line at or above d’=1.9 in Figure 3), an ability that subtly improved with increasing SNR [rANOVA: F(1.46, 26.21) = 11.543, p < 0.001, *rt*^2^_*G*_ = 0.18], consistent with our prior work [15]. To reach performance of d’= 1.9 or better, the target tone needed to be at least 380 ms long, as estimated from tone detection thresholds at 0 dB SNR as a function of tone duration in two additional, non-implanted gerbils (see Figure 4 and results of GLM fit in Table 1). Analysis revealed that behavioral sensitivity asymptotes at approximately 500-600 ms.

**Table 1.**
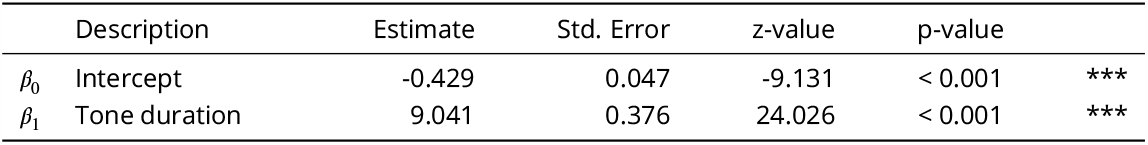
GLM results for behavioral performance vs. tone duration. Degrees of freedom: 143. Model: HR = logit^−1^ (*β*_0_+*β*_1_× Tone duration Significance codes: ‘***’ p < 0.001, ‘**’ p < 0.01, ‘*’ p < 0.05, ‘.’ p < 0.1, ‘ ’ p ≥0.1.

A total of 151 single-units in the freely moving, awake and sound-detecting gerbils showed target-evoked responses in the presence of masking noise, as evaluated offline by visually comparing Go vs. NoGo RTHs. Therefore, no units were excluded from further statistical analysis. First-spike response latencies, shown in Figure 5, were consistent with those typically observed in primary auditory cortex of awake gerbils [27]. During NoGo trials, when only the masking sound was played, RTHs robustly phase-locked to the masker envelope, both in the active and in the passive conditions (light red and light grey lines in Figure 6 A and B track the 10 Hz acoustic envelope of the masker). In the passive conditions, overall firing rate did not appreciably vary across time. However, in the active conditions, the firing rate was modulated by the animal’s behavior, increasing by 21.8% for 200 ms (or two masker cycles) after the onset of the nose poke ([paired t(151) = 6.2, p < 0.001]; note how the black lines are steady until tone onset in Figure 6A, whereas the red lines rise above baseline prior to tone onset, after the nose-poke in Figure 6B), followed by suppression until target tone onset.

**Figure 5.**
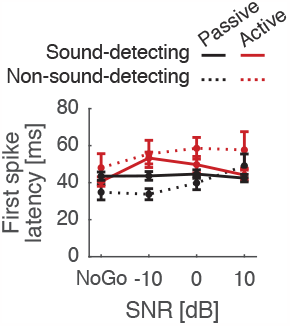
Average first-spike latency of all single-units. Error bars show one SEM.

**Figure 6.**
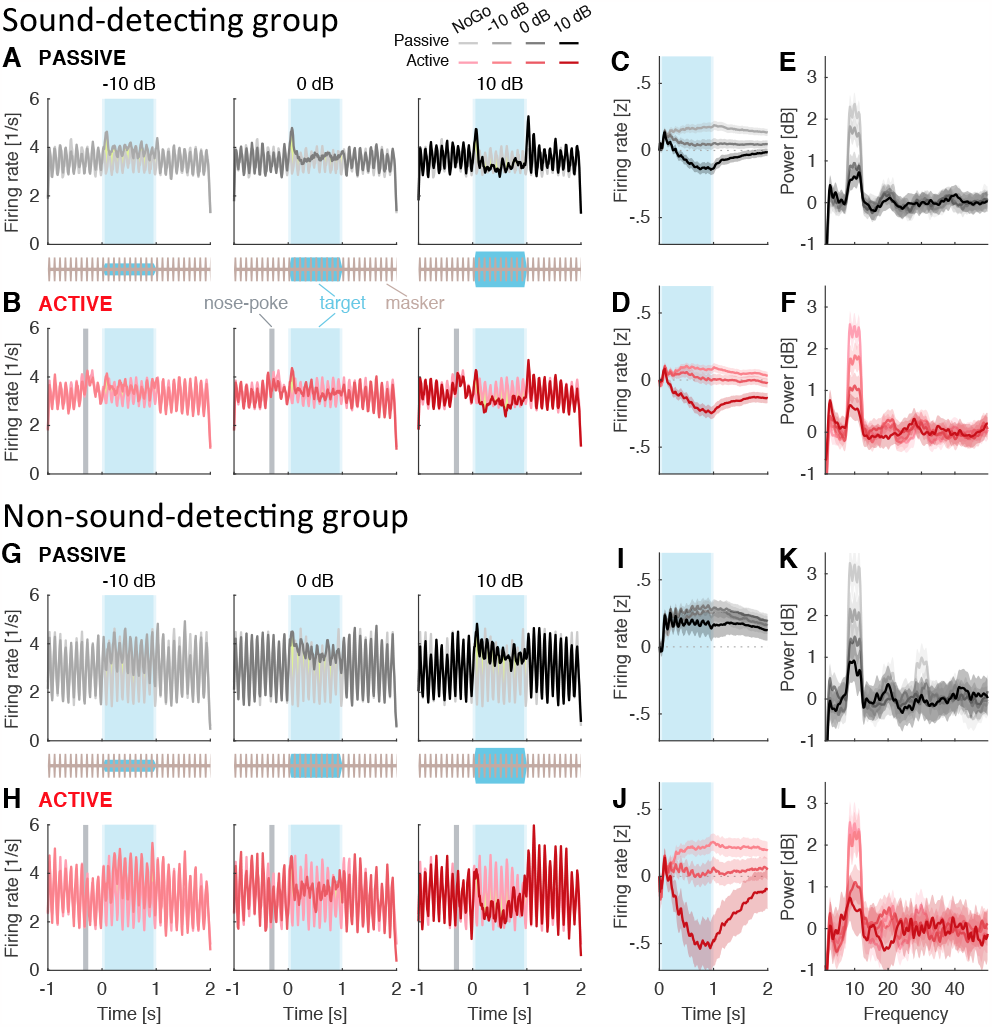
**A**,**B**,**G**,**H** Response time histogram for sound-detecting (n=4, units=151) and non-sound-detecting gerbils (n=2, units=56). NoGo histograms are depicted under each Go histogram for direct visual comparison. Timing of target indicated by blue shading, timing of nose-poke indicated by grey bar in active conditions. **C**,**D**,**I**,**J** Firing rate z-score of the neural response as a function of time, calculated in incremental windows relative to target onset. **E**,**F**,**K**,**L** Power of spectral density of the neural activity calculated with MTS at different frequencies. Ribbons indicate one standard error of the mean (SEM).

In addition, during Go trials, across-unit average RTHs showed target-evoked onset and offset responses (see darker lines around the blue-shaded target time-windows in Figure 6 A and B). Onset enhancement occurred at all SNRs. Offset enhancement was pronounced at 10 dB SNR, but weak or absent at 0 and -10 dB SNR. During the sustained portion of the target sound, from 100 to 950 ms, suppression occurred at 10 dB SNR, vs. enhancement at 0 dB and -10 dB SNR, relative to the response to the masker alone (dark lines fall below zero in Figure 6 C and D, whereas the lighter lines stay positive throughout the sustained time interval). Despite these non-monotonic changes in sustained firing rate between Go and NoGo trials, power spectral density analysis confirmed the strength by which the single-units tracked the masker *monotonically* decreased with increasing SNR (Figure 6 E and F show sharp peaks near 10 Hz, peak height increasing with decreasing SNR). Consistent with this, Figure 7B shows for both active and passive conditions, vector strength decreased monotonically with increasing SNR [rANOVA F(2.4, 350.2) = 53.2, p < 0.001, *rt*^2^_*G*_ = 0.112], with no appreciable differences in vector strengths between active vs. passive listening [rANOVA F(1, 145) = 5.5, p = 0.020, *rt*^2^_*G*_ = 0.004]. This is consistent with envelope-locking suppression induced by the target.

**Figure 7.**
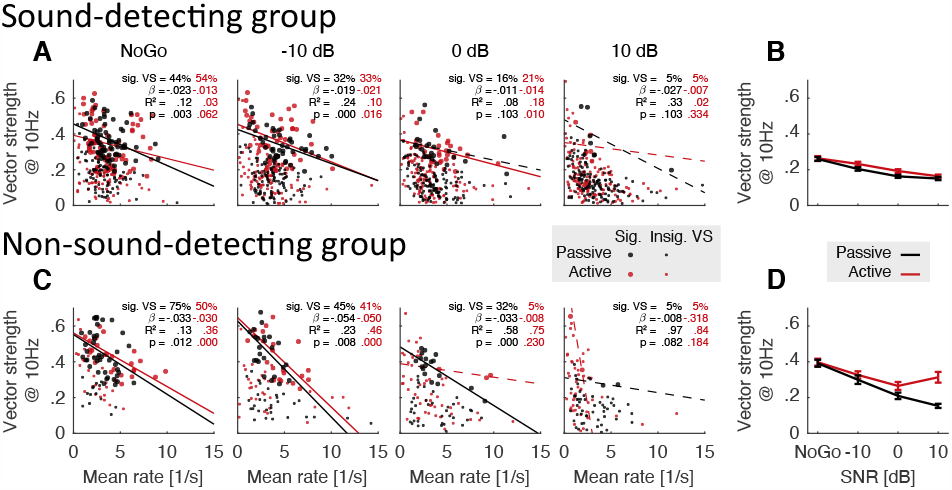
**A**,**C** Vector strength vs. mean firing rate of all units during the sustained response period. Units with a statistically significant vector strength are indicated with larger points. Percentage of these units can be seen at the corner of each plot, along with the details of regression fits over these points. **B**,**D** Vector strength during the sustained response period as a function of SNR. Here only phasic units are included which have at least one significant vector strength across all SNRs and task engagement conditions. Points mark the average, and error bars show one SEM.

There was often a relationship between lower spike rate and stronger envelope locking to the masker when the tone was less likely to be detected. Figure 7A shows the average sustained firing rates vs. vector strengths at 10 Hz for all single-units, with large symbols denoting units with vector strengths that were significantly greater than 0, per Rayleigh’s z-test. Linear regression analysis reveals that at -10 dB SNR and during NoGo trials, units that tended to phase-lock more strongly to the envelope of the masker also tended to have lower firing rates in the passive conditions (notice the near absence of data points with high firing rate and high vector strength in Figure 7A left two panels, solid black regression lines), accounting for approximately 15% of the variance in the data. In contrast, in the passive conditions at 0 dB and 10 dB SNR, and in the active condition at 10 dB SNR, average sustained firing rates and vector strengths were not significantly correlated (dashed regression lines).

Next we wondered whether target-evoked decorrelation, indicated by a decrease in *ρ*, could be a reliable cue for detecting target sound at both positive and negative SNRs. The *ρ* metric is a similarity measure that captures whether temporal information changes when target sound occurs, relative to just having background sound (i.e., Go vs. NoGo). Across both active and passive listening conditions and all SNRs, the target onset sharply reduces *ρ* (Figure 8B, left panel), revealing that in the absence of target energy, single-unit responses are self-similar across trials, and confirming decorrelation as a robust indicator of the target. Note that this analysis includes all single-units. Even single-units with low to absent 10 Hz vector strength, i.e. units that do not appear to track the masker envelope, robustly and repeatably show this trend (see Supplemental Information). In addition, in the active conditions, the sharp reduction in *ρ* is preceded by a small but consistent increase in *ρ* (see time-expanded plot in Figure 8B, right panel), suggesting that after the animal initiates a trial via the nose-poke, the single-unit responses become more sharply tuned to the background sound only, before the addition of target sound decorrelates the temporal structure in their responses relative to just the presence of background sound.

**Figure 8.**
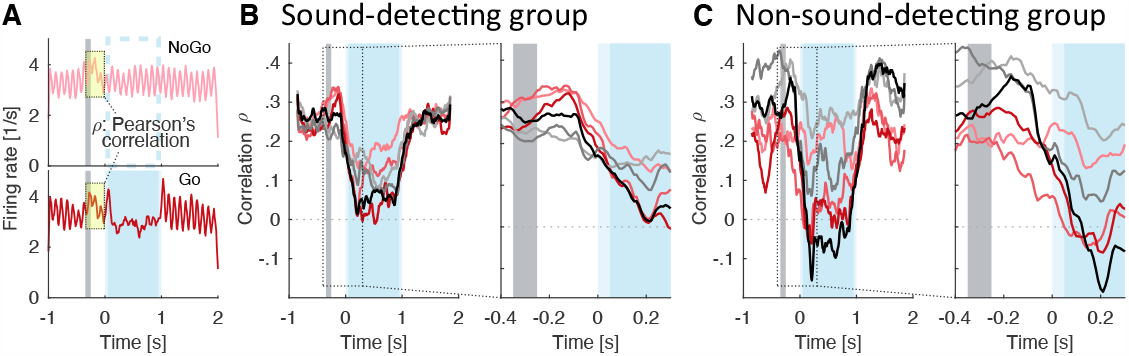
**A** Schematic calculation of Pearson’s correlation *p* of neural responses for a 300 ms time-window starting at nose-poke. Firing rates at matching time points are correlated between each Go condition and the NoGo. **B**,**C** Running Pearson’s correlation of neural responses during Go trials in reference to NoGo. Right panels are time-expanded on the x-axis. Timing of target indicated by blue shading, timing of nose-poke indicated by grey bar.

### Non-Sound-Detecting Gerbils

The passive conditions were always recorded following active testing on the same day, after the animal had reached satiety. One striking observation from the sound-detecting gerbils above is the high similarity in the single-unit responses between active vs. passive conditions (compare Figure 6 A vs. B). This raises the possibility that in the nominally passive listening conditions, these highly trained animals were, in fact, not ignoring the sounds reaching their ears, even though they were not actively seeking rewards. As a control for this caveat, we next tested gerbils that had been conditioned to expect a water reward simply for initiating the nose poke, even when no tone was playing, i.e. both Go and NoGo trials were always rewarded. Note that while normally-hearing gerbils can be trained to reliably detect target sound in the current task even at -20 dB SNR over a dozen training sessions, naive gerbils typically cannot perform the task at negative SNRs [15].

In this non-sound-detecting group, we isolated 56 target-responsive single-units. Visual inspection of the RTHs reveals target-evoked onset or offset responses at 10 dB SNR, but much reduced or absent onset/offset responses at 0 and -10 dB SNR (Figure 6 G and H). In the passive conditions, the presence of target energy consistently increased the firing rate as compared to background-sound-only (all lines fall above zero in Figure 6I). This response pattern was qualitatively different from the sound-detecting group (compare Figure 6C vs. I). Not only were the firing rate differentials between Go and NoGo consistently positive, but they also fluctuated at 10 Hz (note how all lines are positive in Figure 6I, and fluctuate at 10 Hz). In the active conditions, the difference in firing rates between Go and NoGo sustained responses decreased with increasing SNR, a difference that interacted with SNR similar to results from the sound-detecting gerbils (compare Figure 6 J vs. D). Specifically, Go firing rates were suppressed relative to NoGo firing rates at 10 dB SNR (dark red line falls below zero in Figure 6 D and J), but enhanced relative to NoGo firing rates at 0 and -10 dB SNR (lighter red lines fall above zero in Figure 6 J and D). Of note, unlike in the sound-detecting gerbils, firing rates did not increase after the nose poke onset and even slightly decreased by 2.6% [paired t(56) = -3.6, p < 0.001].

Power spectral density was tuned to 10 Hz, and the 10 Hz peak height decreased with increasing SNR, consistent with the idea of tracking the temporal envelope of the masker (Figure 6 K and L). As with sound-detecting gerbils, the relationship between spike rate and envelope locking to the masker was related to tone detectability. In the passive conditions, 10 Hz vector strength and mean firing rate co-varied significantly, except at 10 dB SNR, with lower rate units more likely to significantly phase-lock to the masker envelope (Figure 7C). In the active conditions, this trend was also apparent, but only for NoGo and -10 dB SNR cases, where the non-sound-discriminating gerbils presumably could not hear the target. Similar to the sound-detecting gerbils, here, across-unit average vector strength decreased with increasing SNR in the passive conditions (Figure 7D). However, in the active conditions, vector strength plateaued between 0 and 10 dB SNR. Importantly, overall vector strength was approximately 0.1 higher in the non-sound detecting as compared to the sound-detecting group, indicating weaker envelope-locking suppression induced by the target (compare vertical ranges across Figure 7 B and D, [mANOVA F(1, 200) = 68.2, p < 0.001, *rt*^2^_*G*_ = 0.103]).

Target-evoked response correlations *ρ* were considerably more variable across conditions, as compared to *ρ* in the sound-detecting gerbils (compare vertical spread of curves in Figures 8 B vs. C). However, even in the non-sound-detecting gerbils, *ρ* consistently decreased in the presence of target sound. Of note, unlike for the sound-detecting gerbils, nose poke initiation was not followed by increased *ρ* (red lines do not increase to the right of the grey bar in the time-dilated curve of Figures 8C, right panel).

### Neurometric Analysis

To test our core hypothesis that SNR shapes reliance on short-term temporal processing, we next analyzed how rate vs. temporal coding strategies varied with SNR by comparing NoGo vs. Go responses (Figure 9). Mutual information between NoGo and Go spike probability distributions, shown in Figure 9A, did not appreciably vary with SNR [mANOVA F(1.9, 368.7) = 6.9, p = 0.001, *rt*^2^_*G*_ = 0.009], consistent with the observation that gerbils can be trained to hear the target at all SNRs.

**Figure 9.**
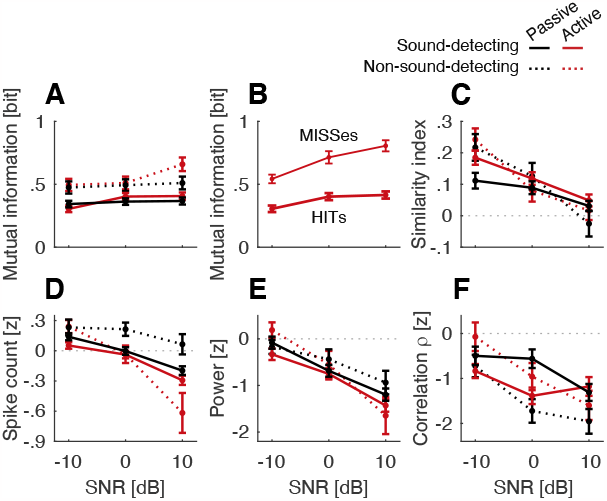
Average of all neurometric rate and temporal measures that were calculated relative to NoGo for each gerbil group and task engagement condition. These metrics were derived for the sustained response period. **A** Mutual information (rate-based measure). **B** Mutual information for the sound-detecting gerbils during active task engagement, separated by HIT and MISS trials. **C** Similarity index calculated as slope of a linear regression fit (combining temporal- and rate-based information). **D** Spike count z-score (rate-based measure). **E** Power spectral density z-score at 10 Hz derived with MTS (temporal measure). **F** Target-evoked correlation response (*p*-index) z-score (temporal measure). Error bars show one SEM.

Surprisingly, sound-discriminating gerbils had much reduced spike rate-based mutual information, as compared to non-sound discriminating gerbils, indicating more similarity between Go and NoGo trials [mANOVA F(1, 191) = 31.6, p < 0.001, *rt*^2^_*G*_ = 0.044]. There was no overall effect of active vs. passive task engagement [mANOVA F(1, 191) = 0.9, p = 0.356, *rt*^2^_*G*_ < 0.001]. However, mANOVA revealed a crossover interaction of SNR and engagement [mANOVA F(2, 382) = 3.1, p = 0.048, *rt*2 *G* = 0.004], consistent with the observation that for both groups of gerbils, the mutual information was somewhat higher during active vs. passive listening at +10 dB SNR.

The association of reduced rate-based mutual information and better task performance was also borne out when separately examining different response types during the Go trials in the active conditions. Figure 9B shows a subset of the data in Figure 9A, specifically: the active sound-detecting data separated into HITs, i.e. Go trials when sound-detecting animals correctly detected the target, vs. MISSes, i.e., Go trials when sound-detecting animals fail to detect a target. The mutual information between NoGo and Go spike distributions during HITs was lower as compared to MISSes [rANOVA F(1, 22) = 45.5, p < 0.001, *rt*2 *G* = 0.268]. The overall effect of SNR remained significant [rANOVA F(2, 44) = 7.8, p = 0.001, *rt*2 *G* = 0.091]. The interaction between mutual information score and SNR was significant, consistent with the observation that at +10 dB, the difference between MISSes and HITs was greatest overall [rANOVA F(2, 44) = 3.5, p = 0.039, *rt*2 *G* = 0.037].

Therefore, the current results show, somewhat paradoxically, that when gerbils were better able to hear the tone, the Go and NoGo responses were more similar to each other based on firing rate as compared to when tone detection performance was worse (HITs vs. MISSes or Sound-Detecting vs. Non-Sound-Detecting gerbils). However, note also that the mutual information metric is agnostic to the readout mechanism employed by the central nervous system. The results are consistent with the interpretation that task training prunes information in the neuronal code, removing neuronal discharge patterns that could be informative because they vary consistently with the presence of the target but that are presumably not helpful to the sensory decoding mechanisms that the animal uses.

Figure 9C shows the similarity index, i.e., the slope of the regression fit linking Go target-background-mixture responses with NoGo responses to the background sound alone [7]. Here, the similarity index did not differ appreciably across groups [mANOVA F(1, 200) = 0.1, p = 0.769, *rt*2 *G* < 0.001], or active/passive listening [mANOVA F(1, 200) = 0.8, p = 0.371, *rt*2 *G* < 0.001], but decreased with increasing SNR across all conditions [mANOVA F(1.8, 366.9) = 45.8, p < 0.001, *rt*2 *G* = 0.061]. This is consistent with the interpretation that the Go -vs. NoGo-evoked responses become increasingly similar with decreasing SNR. The sign of the similarity index is positive across all conditions, suggesting that an SNR-invariant decision metric based on overall similarity could underlie target detection.

The similarity index combines both rate cues and temporal information. To disentangle how these cues drive behavioral task performance across positive and negative SNRs, we next separately analyzed putative decoding mechanisms relying on spike count vs. spike timing. In both groups, the Go-NoGo differences in spike count decreased with increasing SNR (Figure 9D; mANOVA F(1.6, 321.1) = 126.1, p < 0.001, *rt*2 *G* = 0.123), and the magnitude of the Go-NoGo difference was greater in the active than in the passive conditions [mANOVA F(1, 196) = 35.9, p < 0.001, *rt*2 *G* = 0.033]. There was no main effect of group [mANOVA F(1, 196) = 3.0, p = 0.087, *rt*2 *G* = 0.007]. Across all configurations, except for the passive non-sound-detecting configuration, the spike count curve as a function of SNR crossed zero, showing that at -10 dB SNR, more spikes occurred during Go than NoGo trials, whereas at 10 dB, NoGo trials had a higher spike count than Go trials (in Figure 9D, all lines except for dashed black line cross zero). Therefore, a putative neuronal readout relying on Go-NoGo spike count distances would need to use different decision strategies at positive vs. negative SNR, detecting target sound via a decrease in sustained firing at positive SNR, as compared to increased sustained firing at negative SNRs. Also, note that the differences in z-score across SNR were quite small.

In contrast to the spike count, putative temporal readouts varied monotonically with SNR, without crossing zero. Specifically, both the 10 Hz modulation power (Figure 9E) and the *ρ* correlation index (Figure 9F) decreased with increasing SNR [Power: mANOVA F(1.7, 300.7) = 73.4, p < 0.001, *rt*2 *G* = 0.066; *ρ*: mANOVA F(2, 406) = 16.1, p < 0.001, *rt*2 *G* = 0.023], with negative z-distances between NoGo vs. Go at all SNRs. There was no main effect of group [Power: mANOVA F(1, 173) = 0.5, p = 0.499, *rt*2 *G* = 0.001; *ρ*: mANOVA F(1, 203) = 1.2, p = 0.275, *rt*2 *G* = 0.001]. Analyzing the 10-Hz vector-strength across groups reveals that the vector strength was higher in the non-sound detecting as compared to the sound detecting group ([mANOVA F(1, 200) = 68.2, p < 0.001, *rt*2 *G* = 0.103]; vector strength in Figure 7 D was overall higher than in Figure 7B), with a main effect of SNR [mANOVA F(2.5, 505.5) = 79.3, p < 0.001, *rt*2 *G* = 0.121] and main effect of task engagement [mANOVA F(1, 200) = 28.6, p < 0.001, *rt*2 *G* = 0.018]. The Go-NoGo difference in 10 Hz power was greater in the active than passive conditions [mANOVA F(1, 173) = 7.2, p = 0.008, *rt*2 *G* = 0.009], but *ρ* did not differ appreciably between active and passive listening [mANOVA F(1, 203) = .3, p = 0.598, *rt*2 *G* < 0.001].

Together, these results suggest that both rate and temporal cues could be used at the positive SNR. However, at 0 dB SNR, spike count does not appreciably differ between Go and NoGo trials [paired t(413) = -2.6, adjusted p = 0.031, Bonferroni-adjusted for multiple comparisons]. Moreover, at the negative SNR, only decision metrics relying on temporal coding can operate with a decision criterion that is consistent with that at the positive SNR.

## Discussion

Moderate background sound disrupts auditory clarity in people with hearing loss, a challenge that no current clinical treatment approach overcomes for the majority of individuals. Effective comprehension with fluctuating background sound involves “dip-listening” within the energetic dips of the background. Prior behavioral work establishes that both peripheral mechanisms and central auditory processing contribute to dip-listening. Evidence supporting central mechanisms include the comodulation masking release phenomenon, where added masker energy even outside the frequency range of the target can paradoxically improve target detection and identification performance, an effect obliterated by backward-masking, another central phenomenon [28, 29, 30]. Moreover, even up to six months after recovering from an episode of temporary conductive hearing loss from otitis media, normally hearing children experience reduced ability to listen in the dips [31]. Consistent with this, gerbils with chronic juvenile-onset sound deprivation have raised detection thresholds in modulated and unmodulated noise despite not showing widened cochlear filters as compared to normally hearing controls [15, 32, 33]. Even a theoretically ideal hearing aid could not compensate for peripheral dysfunction. In contrast, central processes could be targeted by rehabilitative technology. To better understand how hearing loss disrupts auditory clarity in situations with background sound and how auditory clarity could be restored, we therefore need to define central mechanisms for hearing in background sound. The current study intends to contribute towards this goal by examining SNR invariance in auditory cortex.

To elucidate the factors causing dip-listening to operate across a range of negative and positive SNRs, this study sought to test the hypothesis that listening at negative SNRs is mediated by a stronger reliance on temporal cues, as compared to positive SNRs. Evidence supporting this hypothesis comes from behavioral studies in human listeners showing that an individual’s ability to resolve temporal fine structure information predicts their ability to suppress temporally fluctuating background sound [2, 34].

We here recorded auditory cortical responses in normally hearing, awake, freely moving adult gerbils, while these gerbils either actively detected a target tone in 10 Hz modulated noise or passively heard the same sounds. Gerbils performed with comparable behavioral sensitivity across three SNRs that were either negative, balanced or positive. In the following sections, we discuss how SNR shapes the potency of rate vs. temporal neuronal coding cues for solving this task.

Using the conceptual framework of classic decision theory, a tacit premise of prior work looking at the effect of SNR on tone detection in modulated noise is that the separation between target and masker along an internal decision variable reduces with decreasing SNR. However, this assumption had not explicitly been tested, raising the possibility that SNR shapes decision variables differently depending on whether they rely on rate or temporal coding.

### Spike Count

Spike count differences between Go and NoGo trials tended to be negative at positive SNRs, and positive at negative SNRs (Figure 9D), supporting the idea that a simple decision strategy based on spike count would not suffice to account for the nearly SNR-invariant behavior observed over the tested SNRs in the current work (Figure 3). Intriguingly, in the passive conditions of the non-sound-detecting, we do not see this pattern of SNR-dependent suppression/enhancement reversal (Figures 9D and 6I). Instead, the response time histograms strongly resemble the waveform of the acoustic mixture (Figure 6G), as opposed to a pre-processed trace that signals the presence of a new object. This is consistent with the interpretation that the non-sound-detecting gerbils may not have perceived the tone as a separate object in the passive condition.

### Envelope Locking Suppression

In anesthetized untrained cats, phase-locking to a moderately slow amplitude-modulated masker envelope is suppressed 75 ms after onset of a target tone [7]. This phenomenon, referred to as envelope locking suppression, emerges at the medial geniculate body, sculpting response patterns in primary auditory cortex across a range of positive to negative SNRs, and even for target tones below the quiet threshold of the neurons [7].

The existence of envelope locking suppression was confirmed in single units in the primary auditory cortex of anesthetized and untrained rats [35] as well as mice, albeit with low prevalence across recorded sites [12]. Specifically, in rat, at 0 dB SNR, average neuronal responses to unmodulated masking noise did not change much upon the addition of a tone, but when the masking noise was modulated, the addition of a tone caused suppressed envelope following responses [35]. Furthermore, in mouse, for SNRs ranging from -10 to 20 dB, single units responded more strongly to tones in modulated narrowband noise when this narrowband noise was presented with a coherently or incoherently modulated flanking noise, and showed overall stronger envelope locking suppression in the presence of forward-masking fringes [12]. Moreover, in mouse, with decreasing SNR the firing rate in most units increased, and increased most steeply in the presence of coherently modulated background sound [12]. In contrast to these mammalian responses, neuronal responses in the primary auditory cortex homologue in birds, area L2 of the avian brain, lack locking suppression [36, 37], and have previously been compared to more strongly resemble inferior colliculus responses in cat rather than auditory cortex [7].

We here confirm envelope locking disruption at negative and positive SNRs, both quantitatively through the similarity index (Figure 9C) as well as qualitatively through visual inspection of the response time histograms (Figure 6). However, unlike in the prior work in anesthetized animals, in the current design with awake animals, overall neuronal discharge rates are enhanced, as opposed to suppressed, at negative SNRs (Figure 9D). This phenomenon was observed in all animals that were engaged in a behavioral task, even when those animals were not engaged in sound detection. As a result, spike count differences between Go and NoGo trials tend to be negative at positive SNRs, and positive at negative SNRs (Figure 9D), an observation that is inconsistent with the interpretation that rate coding is SNR-invariant. In contrast, neurometric measures that are based on temporal patterns, including the similarity index and the correlation index *ρ*, do not change signs with decreasing SNR (Figure 9 C and F), making them viable candidate metrics for the behaviorally observed SNR-invariant detection of target sound, across the range of tested SNRs.

Previously, in trained rhesus monkeys that were engaged in a tone detection task with amplitude-modulated background noise, primary auditory cortex recordings showed that single-neuron responses were in-suffcient and pooling of spiking activity across the population of single neurons was necessary to predict behavioral performance from neuronal discharge rates [13]. The monkeys were tested across a range from -5 to 20 dB SNR. However, performance at 0 and -5 dB SNR was at chance, suggesting that the monkeys could not detect the target sound at the two lowest SNRs [13]. Population responses showed enhancement followed by suppression after the target onset, a phenomenon confirmed by the current results for 0 and 10 dB SNR (Figure 6). Our current work supports these findings but shows that at negative SNRs and at the population level, spike count is not a viable metric for tone detection.

In addition, prior work on zebra finches finds evidence for SNR-invariant target feature detectors in primary auditory cortex. When listening for a conspecific target vocalization while a chorus of different conspecific vocalizations was concurrently present in the background, some single auditory cortical neurons selectively encoded the target sound in a manner that is invariant to the background, when those target sounds were behaviorally recognizable [8]. However, the zebra finches in this study could not perform the task at the only tested negative SNR of -15 dB [8]. It is therefore unclear whether the background-invariant encoding of target sound generalizes to negative SNRs. Here, at the single-unit level, only in the sound-detecting gerbils did we observe six highly specialized units with responses in background sound that closely resembled the responses to the target in quiet, across all SNRs (see Supplements FR+/VS+), hinting that the previously proposed selective target encoding mechanism may generalize to the mammalian auditory cortex.

### Task Engagement

In this study, task engagement shaped the anticipatory neural response prior to the potential target onset, as others have reported [17]. Specifically, in sound-detecting animals, the average discharge rate during NoGos did not vary much between active vs. passive listening, except that after the animal initiated a trial via the nose poke, the majority of units showed a characteristic increase in firing rate, followed by a decrease (Figure 6)B. However, in non-sound-detecting animals, firing rate did not increase and even modestly declined after nose poking triggered a trial (Figure 6)H. This suggests that the increase in firing rate immediately after nose-poking is an anticipatory auditory-specific response to support the ability to hear out the target, but only when a gerbil is trying to hear target sound.

### Task Training

Overall, sound-detecting animals had higher non target-evoked discharge rates as compared to non-sound-detecting animals. We observed that the rate-based mutual information between sustained NoGo and Go responses was higher (indicating greater discriminability) for non-sound-detecting than for sound-detecting animals (Figure 9A), and higher when sound-detecting animals missed a Go trial vs. when they responded correctly (Figure 9B). This is consistent with the interpretation that sensory task training reduced the amount of shared information between Go and NoGo responses. While counterintuitive, this may arise in highly trained animals. Animals well-trained on a vowel discrimination task displayed lower firing rate and cortical decoding scores than naive animals [38], and animals well-trained to detect a 1 kHz tone displayed lower cortical discrimination scores for trained vs. untrained frequencies [16]. This reduction in mutual information was not borne out in the other neurometric analyses, where differentiation in neural responses between Go and NoGo stimuli appeared comparable for sound-detecting gerbils vs. non-sound detecting gerbils, for both rate and temporal coding metrics (Figure 9C-F).

One explanation is that mutual information may be reduced if ACx activity is broadly suppressed at the population level during task engagement, as suggested by converging evidence [39, 40, 41, 42]. A subset of target-tracking neurons (6 of 151; see Supplements FR+/VS+) are highly sensitive to the target, such that their masked response at all SNRs is comparable to their response to a robust 60 dB SPL target in quiet -resembling ‘grandmother’ neurons. While the current experiment is not designed to determine which neurons drive the animal’s decision, it is notable that at the population level, VS in sound-detecting animals is reduced at all SNRs compared with non-sound detecting animals (Figure 7B vs D), indicating weakened tracking of the masker. This is consistent with the interpretation that ACx responses to the masker are chronically suppressed in the trained animals. Given that the majority of the units were tracking the masker (Supplemental Information, Figure 2), shutting down that majority by task engagement should reduce mutual information between Go and NoGo trials, allowing the target-enhancing units to contribute more strongly to downstream targets, potentially influencing decision-making and signal detection. Further, it suggests that these highly trained animals have the ability to suppress their cortical response to the ongoing masker, even while not engaged in the task.

### Clinical Implications

Most hearing-impaired individuals whose hearing has been restored via hearing aids or cochlear implants find it difficult to dissociate target sound that they are trying to hear from background sound, a phenomenon called masking. Moreover, even individuals with comparable peripheral audiometric thresholds can vastly differ in how well they can identify masked speech, a phenomenon that is thought to arise from central processing deficits [43, 44, 45, 46, 47]. An extensive literature demonstrates that the ability to listen in the dips is disrupted in individuals with hearing loss and in cochlear implant users [48, 49], suggesting that dip-listening mechanisms could be a key to restoring auditory clarity in background sound for people with hearing loss. Provision of visual lip-reading cues or reduction in target set size can restore a hearing impaired person’s ability to identify target speech at negative SNRs [9, 50]. This phenomenon can be interpreted to mean that auditory dip-listening mechanisms in hearing impaired individuals function normally, provided that the overall *peripheral* SNR is low enough for dip-listening benefits to be effective. However, an alternative interpretation of this work is that provision of temporal cues via lipreading or increased stimulus redundancy via smaller set sizes substitutes malfunctioning auditory temporal cues, filling in *central* access to temporal information that is needed to listen in the dips. We currently lack physiological data to disambiguate these possibilities. The current work demonstrates that both rate and temporal cues can be effective at positive SNRs, but that reliance on temporal information is needed for SNR-invariant hearing at negative SNRs. Moreover, we previously demonstrated that gerbils raised with sound deprivation have reduced dip-listening abilities despite the fact that their peripheral tuning appears intact [15, 32], suggesting that central processes play a key role in dip-listening. Future work is needed to understand whether the temporal information needed for dip-listening at negative SNRs can be augmented through other sensory modalities.

## Acknowledgments

We thank Drs. Todd Mowery, Dan Sanes, Brad Buran, Justin Yao, and Kristina Penikis for many helpful comments and extensive guidance on data acquisition and interpretation. This work was supported by NIH-NIDCD R03 DC014008-01 to AI, NIH-NIDCD R01 DC019126-01 to AI, and NIH-NIDCD R01 DC013314 to MJR.

## Author Contributions

AI conceived the idea. AI, NA, and MJR wrote the main manuscript text. NA carried out the numerical calculations and experimental measurements. NA and AI analyzed and discussed the results. MJR reviewed the manuscript and discussed the results. All authors contributed to scientific discussions about and critical revisions of the article.

## Data Availability

The data that support the findings of this study are available from the corresponding author, AI, upon reasonable request.

