## Supplemental Materials for "Increased reliance on temporal coding when target sound is softer than the background"

### Single-unit response types

The data analysis pipeline is described in Materials and Methods, and is depicted in Figure 1.

In order to identify different patterns of single-unit activity, we categorized units as phasic or tonic based on their ability to follow the envelope of the masker. Pooling across Go and NoGo trials during either passive or active listening, units that had at least one significant vector strength (Rayleigh's $p < 0.001$, see Equation 4 in the main text for details) at the 10 Hz modulation rate of the masker were selected as phasic. The remaining units that were not able to phase-lock to the masker at any of the stimulus or task engagement conditions were categorized as tonic.

Among phasic units, we observed four phenotypes based on changes in firing rate and vector strength in response to the target stimulus during active engagement. This change was measured by comparing average activity of the unit across all trials (Go and NoGo) prior to tone onset, with the response of the unit to 10 dB SNR tones. Single-units that increased in both firing rate and vector strength were labeled as FR+/VS+, units that only had an enhanced firing rate but not vector strength as FR+/VS-, units that suppressed their overall firing rate but were better able to phase-lock to the masker as FR-/VS+, and units that decreased in both firing rate and vector strength were labeled FR-/VS-. These four phenotypes cover all quadrants of a rate/temporal response space.

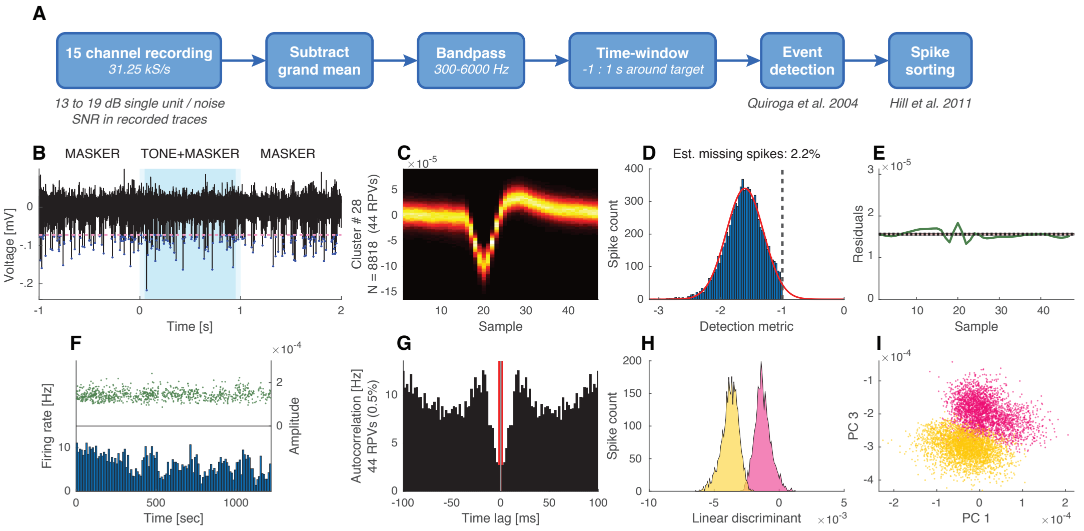


**Figure 1. A** Recorded trace analysis pipeline. **B** A sample of a time-windowed recorded trace. Detection threshold is shown with a pink dashed line, and detected events are marked with blue points. **C-G** Metrics of a representative cluster after automatic spike sorting and visual inspection. **H-I** Separation of two sample clusters with linear discriminant and in principle component space.

| Gerbil group | FR-/VS+ | FR+/VS+ | FR-/VS- | FR+/VS- | Tonic | Total |
|---|---|---|---|---|---|---|
| Sound-detecting | 28 | 6 | 75 | 28 | 14 | 151 |
| Non-sound-detecting | 18 | 1 | 30 | 4 | 3 | 56 |

**Table 1.** Detected unit counts per phenotype for each of the gerbil groups.

### Decoding sensory information by single-unit response type

Table 1 shows unit counts for all 4 phasic phenotypes and also tonics that were detected in each gerbil group. Results show that more than 90% of single-units in both gerbil groups had the ability to follow the envelope of the masker. Among them, the most common phenotype was FR-/VS- with a preference for suppressing both their overall firing rate, and also their phase-locking to the masker in response to the target stimulus.

In all types of phasic and tonic units, sensory information at the population level could predict behavioral sensitivity (Figure 4). For both FR-/VS+ and FR-/VS-, units at 0 and 10 dB SNR average firing rates decreased in Go vs NoGo trials, consistent with envelope locking suppression (Figure 4 A and C). However, at -10 dB SNR, firing rate increased during Go trials as compared to NoGo, a phenomenon not predicted by envelope locking suppression. In quiet, FR-/VS+ units showed suppression of non-sound evoked discharge at the highest sound level, vs. an increase in firing rate at the lowest sound level.

The FR+/VS+ units responded to the tone-masker mixture with a sustained enhancement in firing rate (Figure 2B). Interestingly, this activity pattern was consistent across all SNRs in both active and passive tasks, and closely resembled their response to 60 dB SPL tones in quiet. The spike count z-scores in Figure 4B also reflect this SNR-invariant response strategy, which would be ideal for a masker-agnostic readout of the target stimulus. Sound-detecting animals reveal 6 units out of 151 that follow this pattern.

### Discussion of single-unit response types

The temporal pattern of envelope locking suppression varies non-linearly across units and SNRs. For instance, while a target sound is present, the majority of units, FR-/VS- units, show decreased vector strength in following the masker envelope, a weak target onset response that does not vary appreciably across SNR, and increased sustained firing with decreasing SNR. In contrast, FR-/VS+ units show increased vector strength to the masker envelope, increased target onset responses, and decreased target sustained responses with decreasing SNR. However, for both types of units, decorrelation between NoGo and Go is a robust, SNR-invariant cue.

When the target energy dominated the acoustic mixture, at 10 dB SNR, the majority of single-units during active listening decreased their firing rate (FR-) as compared to when only masker energy was present, in both sound-detecting (103/151) and non-sound-detecting (48/56) animals. Of those rate-suppressing units, 73% in sound-detecting and 63% in non-sound-detecting animals also decreased the vector strength (VS-) by which they tracked the masker. However, these nominally rate-suppressing units also typically showed a small but consistent *enhancement* in firing rate at -10 dB SNR, rejecting the idea of an SNR-invariant rate-based decision criterion from these units. Intriguingly, unlike in the active modes, and unlike in sound-detecting animals, in the passive responses of non-sound-detecting animals, firing rate consistently increased after target onset (Figure 9 G and I in the main text). Rate-based neurometric sensitivity in rate-enhancing (FR+) units of non-sound-detecting and passively listening animals did not vary appreciably was positive at all SNRs.

In contrast, across all rate-suppressing units, in sound-detecting animals, decorrelation $d'$, a template model based on the magnitude of the decorrelation of the signal reaching the ears prior vs. during tone onset, is directly proportional to behavioral sensitivity and does not vary appreciably with SNR. In non-sound-detecting animals, the decorrelation $d'$ analysis predicts an ability to detect the target at 10 and 0 dB, but not -10 dB. This prediction mirrors the learning curves in the behavioral data demonstrating that at the beginning of training, gerbils cannot detect a tone at -10 dB SNR. Indeed, it often takes more than 20 training sessions until gerbils are reliably able to signal the presence of a tone at -10 dB SNR.

When responding to a target in quiet, during passive listening, the RTHs of the populations of rate-suppressing units tended to display strong onset responses followed by sustained suppression. In contrast, rate-enhancing unit responded more slowly to target sound and in sustained patterns. FR+/VS+ units also showed offset responses. This hints that these units rebound after inhibition in the presence of target sound [1]. Intriguingly, in the presence of target sound, FR+/VS+ units increase the vector strength by which they track the masker. However, despite this increased envelope locking, decorrelation-based neurometric sensitivity decreases with increasing SNR, and is small or absent at 0 and 10 dB SNR. In contrast, rate-based neurometric sensitivity appears to be roughly SNR-invariant, at least over the range of tested SNRs. Indeed, unlike in the other three unit types, the RTH patterns of FR+/VS+ unit are comparable across SNRs, but differ from their responses to the same targets in the absence of background sound.

Together, these findings show that at the level of core auditory cortex, at least two readout mechanisms exist. The majority of units change their firing rate when target sound is added. In non-sound-detecting and passively listening animals, firing rate increased at all SNR. In sound-detecting animals or in tasked-engaged non-sound-detecting animals, firing rate depended non-monotonically on SNR. Although, decorrelation in the envelope locking response of these units, either through a decrease or an increase in envelope locking, varies consistently with a change in behavioral sensitivity and is robust to SNR.

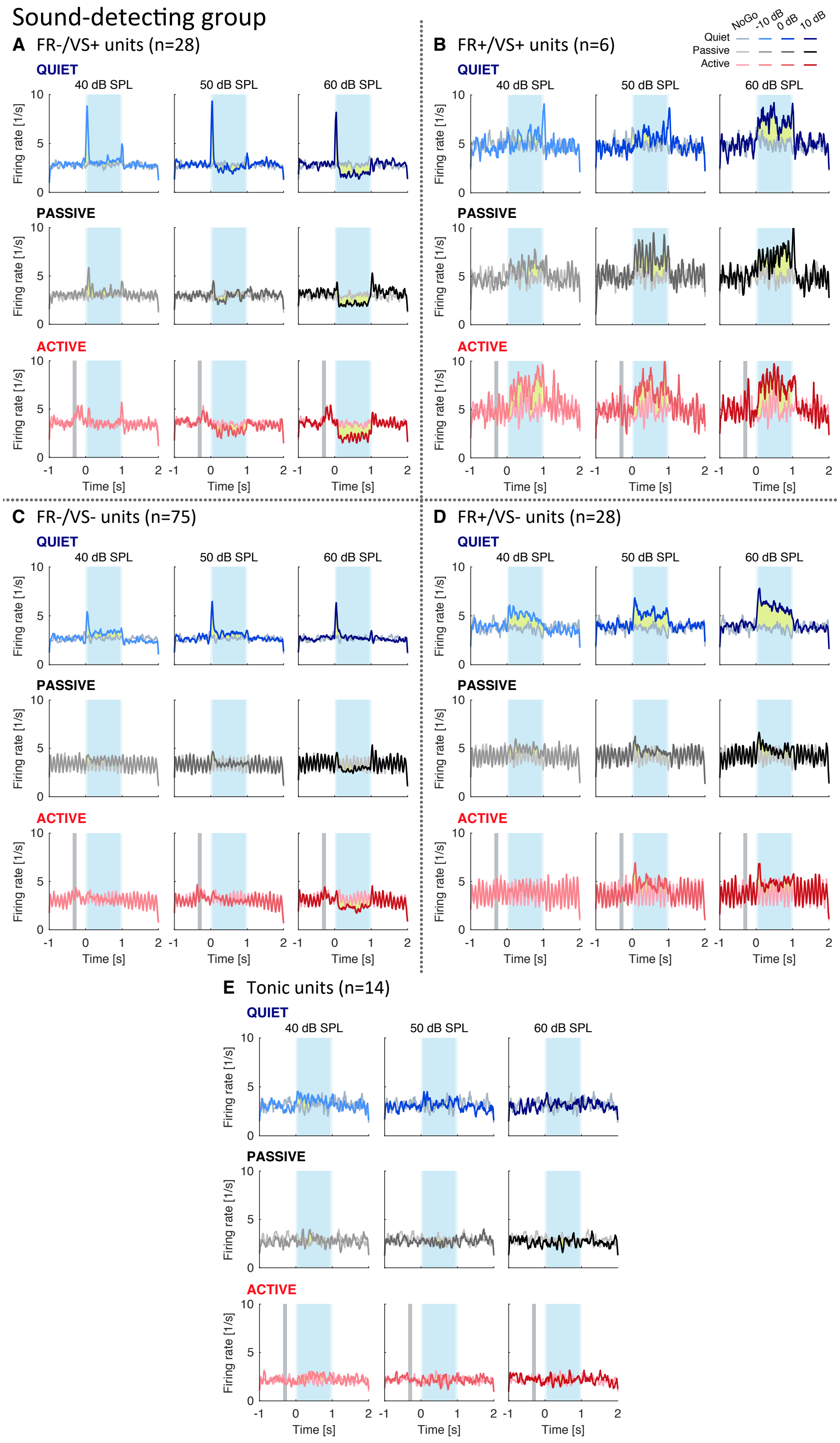


**Figure 2.** RTH of different unit phenotypes for sound-detecting gerbils. In addition to the passive and active engagement conditions with modulated masker (shades of black and red, same as Figure 6 in the main text), here, unit responses to pure tone in quiet (i.e. in absence of background sound) are also illustrated as shades of blue.

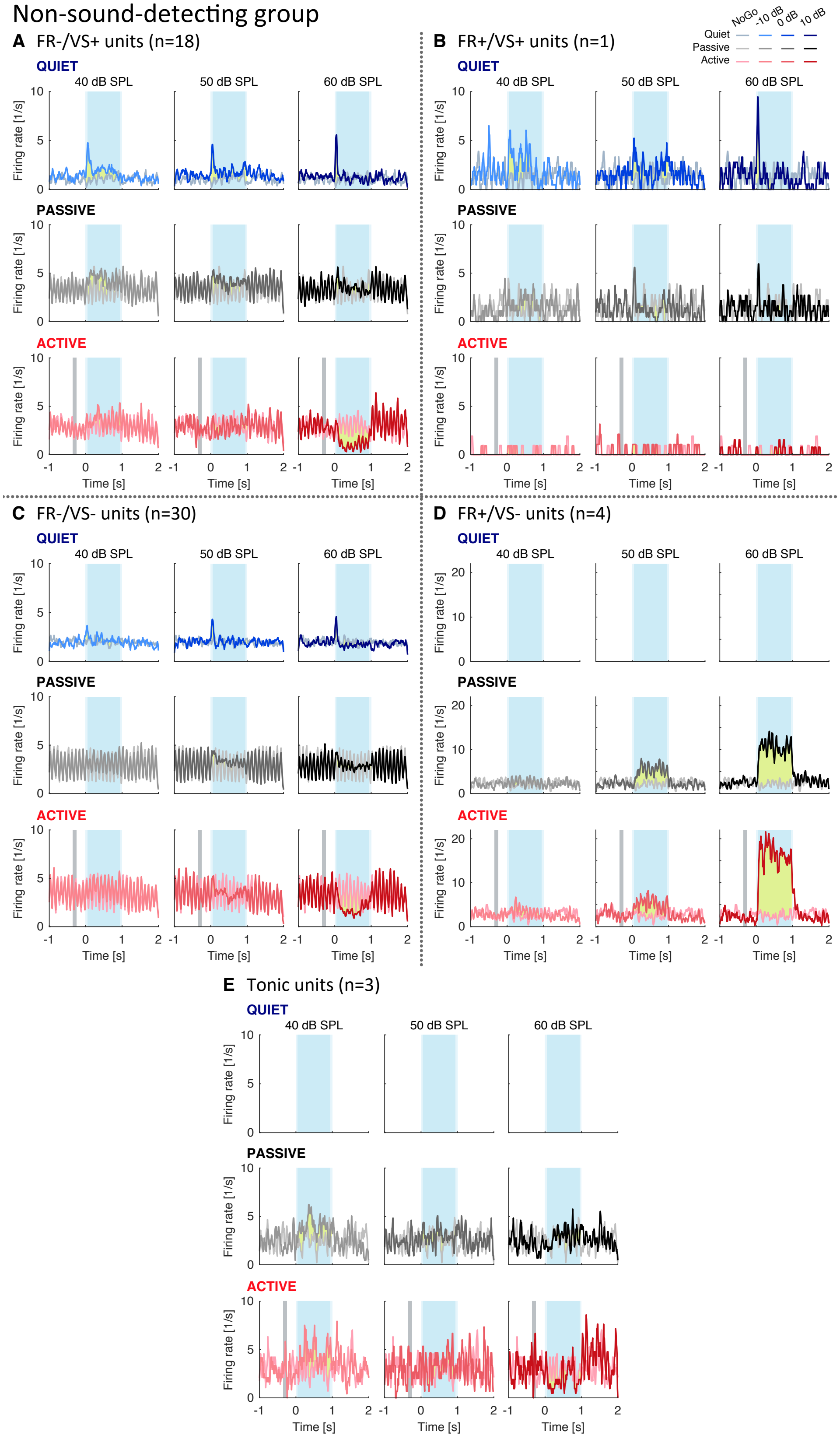


**Figure 3.** RTH of different unit phenotypes (similar to Figure 2) for non-sound-detecting gerbils. Note: FR+/VS- and tonic units have missing recordings for the quiet condition.

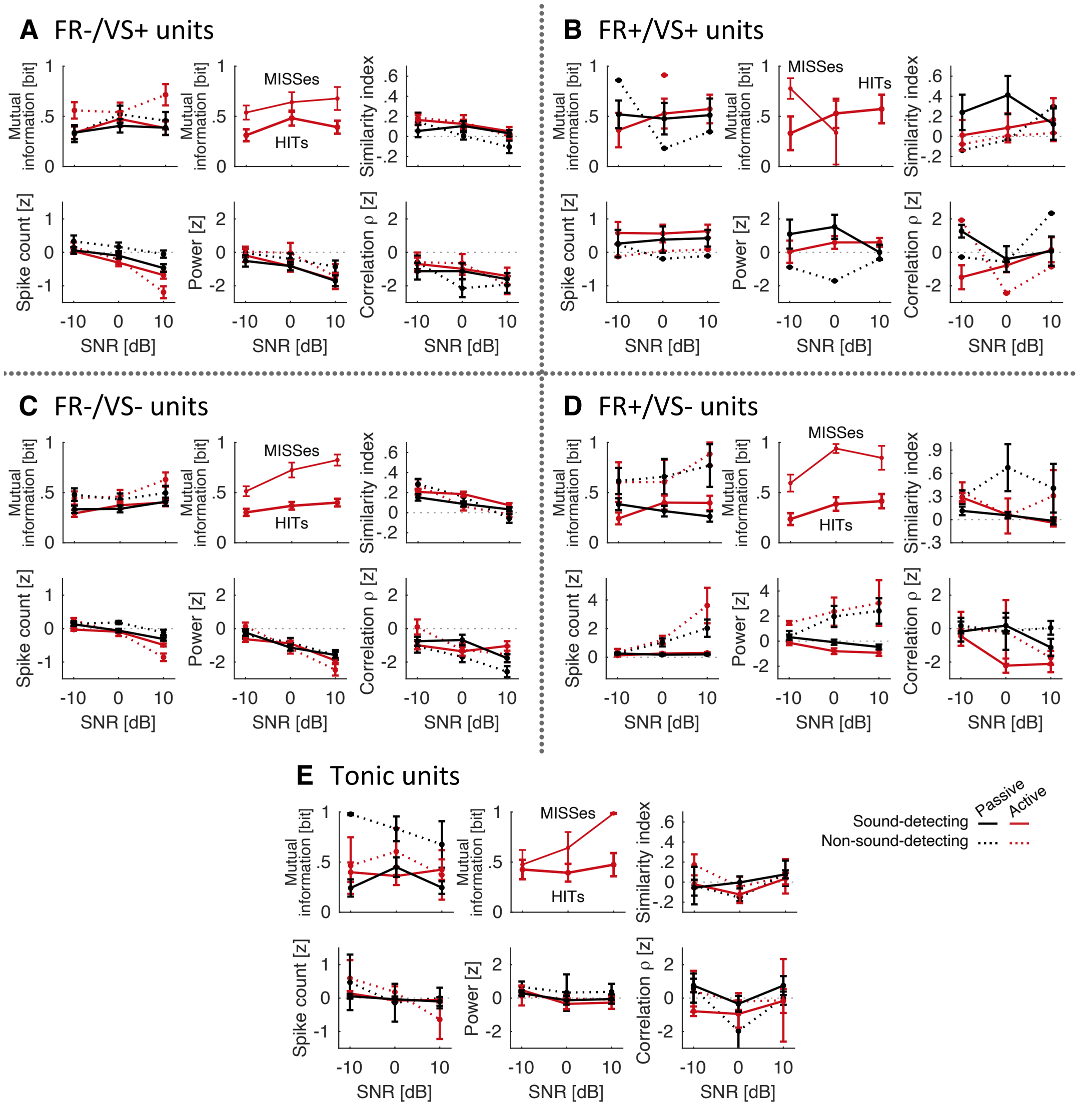


**Figure 4.** Average of neurometric rate and temporal metrics (similar to Figure 9 in the main text) broken down per each of the five unit phenotypes. For unit counts per phenotype for each gerbil group, refer to Table 1.
